# Polymerase and primer biases in eukaryote ultra-long-read metabarcoding

**DOI:** 10.64898/2026.11.04.764338

**Authors:** Leho Tedersoo, Valeria Flores-Almaraz, Rasmus Puusepp, Vladimir Mikryukov

## Abstract

Ultra-long-read metabarcoding is a recently developed method for biodiversity monitoring that enables both high-accuracy species-level identification and reliable phylogenetic placement of taxa. Ultra-long-read analyses require high-processivity DNA polymerases for amplicons and primer pairs that fully cover the targeted taxonomic groups. So far, there is no published information on the relative suitability of commercial high-processivity polymerase kits and on the relative performance of primer pairs targeting a DNA fragment spanning much of the 18S, ITS and 28S rRNA gene regions in eukaryotes. Using PacBio sequencing, we assessed the performance of five commercial high-processivity DNA polymerases and three ultra-long marker primer pairs relative to a commonly used full-ITS primer pair on producing artefacts and recovering biodiversity in two mock communities and 10 soil samples. Our analyses revealed great differences in both polymerases and primer pairs in data quality and biodiversity. For testing evolutionary hypotheses, we recommend using the Solis or Fusion polymerases and the Euk575F-21Rngs primer pair in ultra-long-read metabarcoding analyses of soil eukaryotes. Compared with a previous study using the same DNA samples, the ultra-long amplicons revealed substantially lower biodiversity than medium-sized, full-ITS amplicons across taxonomic levels, suggesting that medium-sized amplicons may be better suited for routine soil eukaryote biodiversity monitoring than ultra-long amplicons.

## 1. Introduction

Metabarcoding of environmental DNA (eDNA) has become a key technique for monitoring biodiversity of microorganisms (Franzosa et al. 2015) and certain macroorganisms, such as fish (Miya 2022), arthropods (Elbrecht et al. 2019) and higher plants (Deiner et al. 2017). The so-called second-generation, short-read sequencing platforms, such as those from Illumina, are the most widely used for metabarcoding due to their low data costs and ease of data analysis (Deiner et al. 2017). Yet, the short reads they produce commonly lack taxonomic and phylogenetic resolution for accurate species discrimination (Tedersoo et al. 2021).

The third-generation, long-read sequencing methods, PacBio and Oxford Nanopore, have been on the market for nearly a decade (Gonzalez et al. 2025). However, only recent advances in sequencing chemistry and basecalling enable the sequencing of ultra-long reads (>3 kb amplicons) at high accuracy and at a reasonable cost (Bista & Lino 2026). Most such long-read studies focus on much of the rRNA gene locus, including the small subunit (SSU; 16S or 18S), the Internal Transcribed Spacer (ITS) region and large subunit (LSU; 23S or 28S). At least in theory, variable regions, such as ITS, SSU V4-V5 subregions and LSU D2-D3 subregions, offer ample opportunities for species- and genus-level identification, while more conserved subregions enable accurate phylogenetic placement (Tedersoo et al. 2021; Jamy et al. 2022). For microeukaryotes, ultra-long reads can also be used to discover and describe novel taxa (Tedersoo et al. 2026a).

Since only a few tens of published studies have used ultra-long reads for biodiversity analyses, there is significant uncertainty about the knowledge-based selection of optimal methods, including primer, polymerase, quality-filtering and identification strategies (Gonzalez et al. 2025; Bista & Lino 2026). Studies on short-read data show that DNA polymerase choice influences error rates, chimera formation and community representation (Gohl et al. 2016; Sze & Schloss 2019; Li et al. 2020; Nagai et al. 2021), primarily depending on the polymerase proofreading activity (Oliver et al. 2015), template AT/GC ratio, presence of long homopolymers (Nichols et al. 2018) and PCR conditions (Brandariz-Fontes et al. 2015). Short-read and full-ITS amplicon studies demonstrate the importance of primer choice for biodiversity recovery in eukaryotes (del Campo et al. 2019; Tedersoo & Anslan 2019; Li et al. 2020; Owens et al. 2024), mainly through primer-template matching (Tedersoo et al. 2015; de Filippis et al. 2017). Other factors that influence the accuracy of biodiversity estimates include PCR cycle numbers (Haas et al. 2011; Gohl et al. 2016) and selection of marker subregion (Tedersoo et al. 2015; Li et al. 2020).

Using PacBio metabarcoding of ultra-long amplicons, we tested the relative performance of three primer pairs and five commercial, high-processivity PCR kits on data quality and biodiversity recovery in two mock communities and ten composite soil samples (see Figure 1 for design). We focused on the following research questions: i) what are the best suitable PCR kits and polymerases; and ii) what are the most suitable primer pairs. By using comparable sequence data from mock communities and soil samples in previous publications, we also evaluated the pros and cons of ultra-long amplicons (3-6 kb) compared with shorter full-ITS amplicons (0.5-1.5 kb).

**Figure 1.**
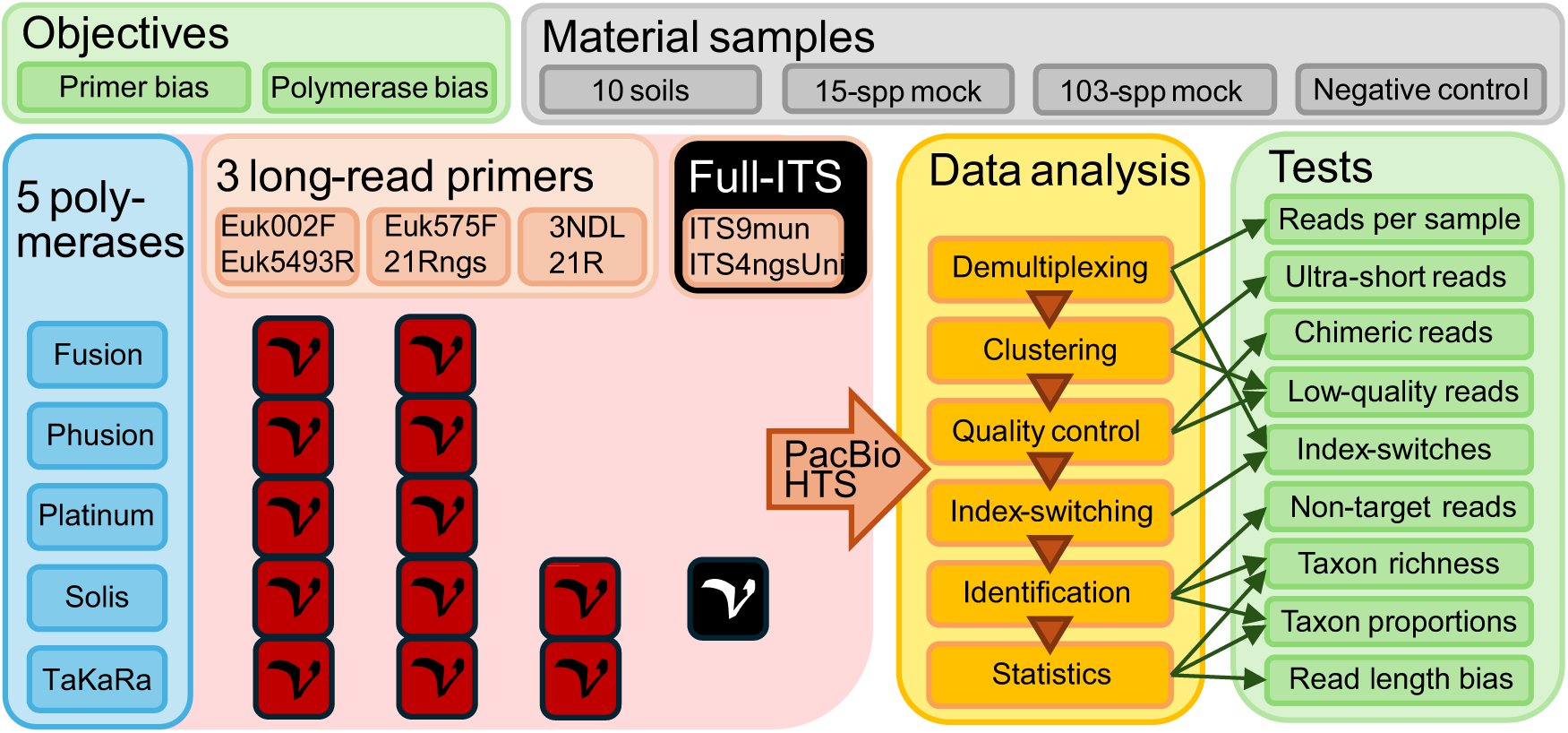
Conceptual overview of the layout of the study, including asymmetric primer pair and DNA polymerase kit combinations. Black-shaded primers and combinations indicate the data borrowed from previous studies.

## 2. Methods

### 2.1. Materials used for analyses

We compiled two mock communities based on DNA extracted from 103 fungal fruit-bodies (Tedersoo et al. 2026b) or 15 fungal cultures (Table 1). Fungi were preferred over other eukaryotes due to lower expected contamination and our experience with these taxa. The 103-species mock community had low phylogenetic diversity, comprising largely *Cortinarius* and *Russula species*. The 15-species mock community included members from different orders of various phyla. Both mock communities were prepared from equal proportions of DNA mixtures as described in Tedersoo et al. (2026b). Ten composite soil samples from different sources were selected for analysis based on previous successful amplification of 1-kb rRNA gene amplicons and relatively high species and phylum-level eukaryote biodiversity (Table 2). The soil samples were collected and processed as described in Mikryukov et al. (2026b).

**Table 1.** Members of the 15-species mock community.

| Taxon name | Culture ID | Sequence accession* | Country |
|---|---|---|---|
| *Xerocomus chrysenteron* | TFC200500 | UDB01088340 | Estonia |
| *Endomyces scopularum* | TFC202287 | UDB0778304 | Estonia |
| *Inonotus obliquus* | TFC202530 | UDB0801836 | Estonia |
| *Tricholoma stiparophyllum* | TFC200493 | UDB01088336 | Estonia |
| *Amphinema byssoides* | TFC200535 | UDB01088360 | Estonia |
| *Parengyodontium album* | TFC200536 | UDB01088361 | Estonia |
| *Rhizopogon mohelnensis* | TFC200419 | UDB01088325 | Estonia |
| *Armillaria borealis* | TFC200780 | UDB01088369 | Estonia |
| *Rhizopus microsporus* | TFC202680 | UDB07672067 | Estonia |
| *Tomentellopsis submollis* | TFC200423 | UDB01088328 | Estonia |
| *Clitopilus cystidiatus* | TFC200516 | UDB01088350 | Estonia |
| *Cephalotrichum stemonitis* | TFC202694 | LR875174 | Estonia |
| *Ganoderma applanatum* | TFC202745 | UDB07675754 | Estonia |
| *Dacrymyces tortus* | TFC202573 | MW191999 | Estonia |
| *Encoelia furfuracea* | TFC202678 | KX090796 | Estonia |

\* Accession numbers include sequence data deposited in UNITE (“UDB”) or GenBank (others).

**Table 2.** The soil samples used in the analyses. More details are given in Mikryukov et al. (2026b).

| Sample ID | Habitat type | Country |
|---|---|---|
| TUE028516 | flooded grassland | Estonia |
| TUE004987 | flooded grassland | Estonia |
| TUE028662 | salt marsh | Saudi Arabia |
| TUE002021 | broadleaf forest | Dominica |
| TUE001939 | broadleaf forest | Gabon |
| TUE028580 | broadleaf forest | Panama |
| TUE028104 | broadleaf forest | Suriname |
| TUE028307 | broadleaf forest | Indonesia |
| TUE000611 | mangrove | Colombia |
| TUE001883 | woodland | Cuba |

To assess the relative performance of PCR kits and high-processivity polymerases, we selected five kits that were available through local vendors: Herculase II Fusion DNA Polymerase (hereafter Fusion; Agilent Technologies, Santa Clara, CA, USA), Phusion Plus PCR Master Mix (Phusion; Thermo Fisher Scientific, Waltham, MA, USA), Invitrogen Platinum SuperFi II PCR Master Mix (Platinum; Thermo Fisher Scientific, Waltham, MA, USA), Solis Hot Start HiFi Master Mix (Solis; Solis BioDyne, Tartu, Estonia) and TaKaRa Advantage 2 Polymerase Mix (TaKaRa; Takara Bio, Kusatsu, Japan). Previously, we have used the TaKaRa and Solis kits to populate the EUKARYOME long-read reference database (Tedersoo et al. 2024). The TaKaRa polymerase has also been used to assess the performance of ultra-long rRNA gene amplicons (Wurzbacher et al. 2019; Jamy et al. 2020; Fang et al. 2023) and to test phylogenetic diversity hypotheses (Jamy et al. 2022). The Fusion polymerase has been used in a pioneering long-read survey of aquatic fungi (Heeger et al. 2018). The Phusion polymerase has been used for nanopore sequencing of bacterial 16S-ITS-23S amplicons (Cusco et al. 2018; Petrone et al. 2023) and benchmarking eukaryote long-read analyses (Chwalinska et al. 2025). The Platinum polymerase has been used to construct the Microflora Danica environmental microbiome atlas (Overgaard et al. 2024; Singleton et al. 2026) and in annual monitoring of a lake microbiome (Chauvet et al. 2026).

We selected three primer pairs for a comparative analysis of eukaryote biodiversity. The 3NDF (5’-GGCAAGTCTGGTGCCAG-3’; Cavalier-Smith et al. 2009) and 21R (5’-AGAGACGAGGCATTTGGCTAC-3’; Schwelm et al. 2016) primer pair has been used most widely in previous pan-eukaryotic analyses (Jamy et al. 2020; Jamy et al. 2022; Chauvet et al. 2026; Singleton et al. 2026). The second primer pair, Euk575F (5′-ASCYGYGGTAAYWCCAGC-3′; Aslani et al. 2024) - 21Rngs (5’-GACGAGGCATTTGGCTAC-3’; this study) is located a few bases inside the positions of the previous primers. These primers were specifically co-designed to avoid overlapping common intron positions, account for template mismatches by introducing degenerate bases, and equalize primer melting temperatures. This primer pair serves as the main standard for populating the EUKARYOME database with long reads. Similarly, the Euk002F (5’-AYCTGGTTGATYYYGCCA-3’; this study) - Euk5493R (5’-GTCTAAACCCAGCTCACG-3’; this study) primer pair was designed to account for all eukaryotes, but it covers a much longer region, starting from the second base of SSU and covering roughly 90% of LSU (Figure 2). To compare with these three ultra-long amplicons, we used previous soil (Mikryukov et al. 2026b) and mock community (Tedersoo et al. 2026b) full-ITS sequence data of the standard eukaryote primer pair ITS9mun (5’ GTACACACCGCCCGTCG-3’; Tedersoo & Lindahl 2016) - ITS4ngsUni (CCTSCSCTTANTDATATGC; Tedersoo & Lindahl 2016). This primer pair has been used to generate global datasets of soil and aquatic eukaryotes (Mikryukov et al. 2026b; Prins et al. 2026) and to compare eukaryote communities across multiple substrates (Chen et al. 2026). The ITS region was selected as a shared marker for its greater taxonomic resolution and relative ease of chimera detection. The 3NDF-21R, Euk575F-21Rngs, Euk002F-Euk5493R and ITS9mun-ITS4ngsUni primer pairs have an expected amplicon length of around 4.5 kb, 4.5 kb, 5.7 kb and 0.8 kb, respectively. All primers were checked against a pan-eukaryote alignment spanning SSU and LSU. The 3NDF and Euk575F primers had critical mismatches to Microsporidia, Foraminifera and Porosporidae (Apicomplexa). The 3NDF primer had additional critical mismatches to Anaeromoebida, Abeoformidae, Tetramitia and Fornicata, and it spanned a common intron position in Myxogastrea. The 21R and 21Rngs primers had critical mismatches to certain groups in Microsporidia, Foraminifera and Parabasalia. The Euk002F primer had critical mismatches to Microsporidia, Foraminifera and Cyanidiophyta, while its pair Euk5493R had only one central mismatch in a few eukaryotic groups. However, due to poor coverage of the 5’ SSU and 3’ LSU termini among rare eukaryote lineages in public databases, other poorly matching taxonomic groups may exist.

**Figure 2.**
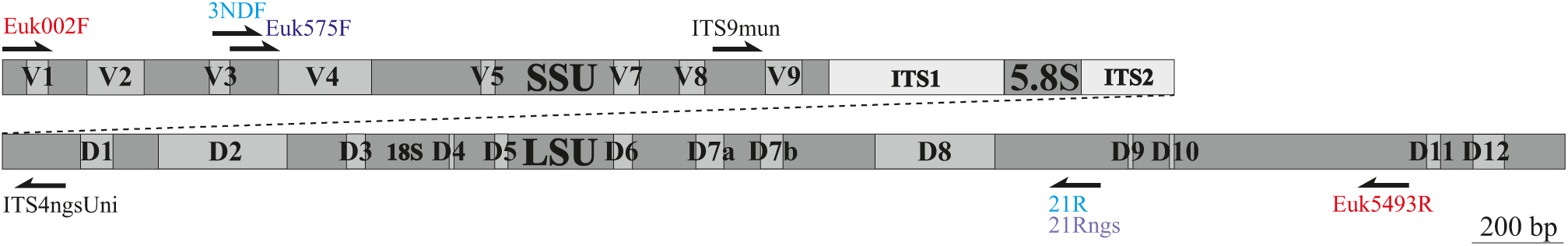
Map of the nuclear rRNA locus indicating variable regions and primers. The variable regions were drawn to scale based on the secondary structure model of *Saccharomyces cerevisiae* (Xie et al. 2012). Colours in primer names indicate the used primer pairs.

All primers were indexed with one of 115, 12-base identifiers that differ from each other by at least 4 bases. The mock communities, negative control samples and soil eDNA samples were amplified in all combinations, except that primer pair 3NDF-21R was used only with Solis polymerase. The ITS9mun-ITS4ngsUni sequences, amplified using Solis 5x HOT FIREPol Blend Master Mix Ready to Load (Solis BioDyne, Tartu, Estonia), were obtained from previous studies using a regular, high-fidelity Solis polymerase (soil samples: Mikryukov et al. 2026b; 103-species mock community: Tedersoo et al. 2026b).

### 2.2. Amplicon preparation and sequencing

The amplification conditions differed between PCR kits and followed the manufacturers’ recommendations. The Fusion polymerase PCRs contained 0.5 µl Herculase II Fusion DNA Polymerase, 5 µl Herculase II Reaction Buffer, 0.31 µl dNTP mix (20 mM each), 0.5 µl forward primer (20 µM), 0.5 µl reverse primer (20 µM), 1 µl DNA extract and 17.2 µl ultra-pure water. The PCR conditions included an initial activation at 95 °C for 2 min, followed by 30 cycles of denaturation at 95 °C for 20 sec, annealing at 60 °C for 20 sec, extension at 72 °C for 150 sec, and a final extension at 72 °C for 3 min. The Phusion polymerase PCRs contained 12.5 µl Phusion Plus PCR Master Mix, 0.5 µl forward primer (20 µM), 0.5 µl reverse primer (20 µM), 1 µl DNA extract and 10.5 µl ultra-pure water. The PCR conditions included an initial activation at 98 °C for 30 sec, followed by 32 cycles of denaturation at 98 °C for 10 sec, annealing at 60 °C for 10 sec, extension at 72 °C for 150 sec, and a final extension at 72 °C for 5 min. The Platinum polymerase PCRs contained 12.5 µl Platinum SuperFi II PCR Master Mix, 0.5 µl forward primer (20 µM), 0.5 µl reverse primer (20 µM), 1 µl DNA extract and 10.5 µl ultra-pure water. PCR conditions included an initial activation at 98 °C for 30 sec, followed by 32 cycles of denaturation at 98 °C for 10 sec, annealing at 60 °C for 10 sec, extension at 72 °C for 150 sec, and a final extension at 72 °C for 5 min. The Solis polymerase PCRs contained 5 µl Solis Hot Start HiFi Master Mix, 0.5 µl forward primer (20 µM), 0.5 µl reverse primer (20 µM), 1 µl DNA extract and 18 µl ultra-pure water. PCR conditions included an initial activation at 95 °C for 15 min, followed by 32 cycles of denaturation at 95 °C for 30 sec, annealing at 62 °C for 30 sec, extension at 72 °C for 4 min, and a final extension at 72 °C for 5 min. The TaKaRa polymerase PCRs contained 0.5 µl TaKaRa Advantage 2 Polymerase Mix, 2.5 µl Advantage 2 PCR Buffer, 0.25 µl dNTP mix (20 mM each), 0.5 µl forward primer (20 µM), 0.5 µl reverse primer (20 µM), 1 µl DNA extract and 19.75 µl ultra-pure water. PCR conditions included an initial activation at 95 °C for 2 min, followed by 35 cycles of denaturation at 95 °C for 30 sec, annealing/extension at 68 °C for 4 min, and a final extension at 68 °C for 7 min. For the Euk002F-Euk5493R primer pair, PCR extension time was prolonged to 3 min for Fusion, Phusion and Platinum polymerases and to 6 min for Solis and TaKaRa polymerases.

The presence of PCR products was checked on 1% agarose gels. Amplicons that produced no or faint bands on the gel were reamplified using 3 additional cycles. PCR products were pooled into two ultra-long-read libraries using 1-20 µl of amplicon per sample based on their band intensity on the gel. Pooled amplicon libraries were cleaned using the FavorPrep GEL/PCR Purification Mini Kit (Favorgen Europe, Vienna, Austria).

PacBio SMRT library preparation and sequencing were performed on a PacBio Revio instrument as described in Tedersoo et al. (2026b). Additional samples were sequenced on the same 25M SMRT cell. The SMRT cell produced 12,861,972 HiFi reads, with a mean polymerase read length of 105,300 bases.

### 2.3. Bioinformatic analyses

Bioinformatic processing was performed with the NextITS pipeline v.1.2.0 (Mikryukov et al. 2026a), a scalable metabarcoding workflow dedicated to ITS sequence analysis, as implemented in Nextflow v.25.10.4 (Di Tommaso et al. 2017). Demultiplexed PacBio HiFi reads were processed separately by primer pair. Quality control was performed with VSEARCH v.2.31.0 (Rognes et al. 2016) and SeqKit v.2.13.0 (Shen et al. 2024). Reads with four or more ambiguous bases, an expected error rate of at least 0.01 or homopolymer stretches longer than 25 bases were removed (Tedersoo et al. 2026b). Primers were trimmed with Cutadapt v.5.2 (Martin 2011), allowing up to two mismatches, and reads lacking either primer were discarded. To compare artefact formation among polymerases and primers, several NextITS filters were disabled: the target region was set to “none”, and Step-1 chimera and tag-jump filtering were excluded. In NextITS Step-2, sequences from all sequencing runs were aggregated and globally dereplicated with VSEARCH using the abundance-sorted clustering algorithm (“--cluster_size”) at 99% sequence similarity. For the ultra-long data subset, the reads were trimmed to the shortest primer sites (Euk575F and 21Rngs) and clustered with VSEARCH as above. The full-ITS data subset was trimmed to the full-length ITS region using ITSx v.1.1.3 (Bengtsson-Palme et al. 2013). Sequence count tables were generated in R v.4.6.0 (R Core Team 2025), using data.table v.1.18.4 (Barrett et al. 2025) and Apache Arrow v.24.0.0 (Richardson et al. 2025).

For taxonomic assignments of OTUs, taxa in the ultra-long data subset were subjected to extraction of the full ITS, SSU and LSU regions from representative sequences using ITSx v.1.1.3. The BLASTn algorithm of BLAST+ v.2.17.0 (Camacho et al. 2009) was used to search 10 best matches separately for the SSU, ITS and LSU regions against EUKARYOME v.2.0. For additional taxonomic and chimera checks, the first 500 bases of SSU and the last 500 bases of LSU were BLASTn-searched.

OTUs were taxonomically assigned primarily based on ITS sequences, using some manual curation as described in Tedersoo et al. (2026b). Protists and animals with limited ITS reference coverage were also evaluated using the SSU V4-V5 subregion. If the OTUs were unassigned to a genus, the best matches of SSU sequences with >99% sequence similarity were used for genus-level assignments. For kingdom-, phylum- and class-level assignments, we used SSU sequence similarity thresholds of 90%, 93% and 95%, respectively.

### 2.4. Read quality assessment

To calculate the error rate of PacBio ultra-long reads for all polymerases and primers, we selected the five most common species in the mock communities for detailed analyses. All reads assigned to the selected OTUs in the mock samples were extracted from the NextITS membership table, along with sample and quality information. The SSU region was delimited with Infernal cmsearch v.1.1.5 (Nawrocki and Eddy 2013) using the Rfam RF01960 covariance model (Ontiveros-Palacios et al. 2024) and the region from the Euk575F primer position to the 3’ terminus of SSU was retained, yielding fragments of roughly 1215 bases. The SSU was selected, because it is the most conserved subregion and was expected to have no intra-individual variation. From each primer, polymerase and species combination, we randomly selected 100 reads with a maximum Phred score (40). Using MAFFT v.7 (Katoh & Standley 2013), we performed separate multiple sequence alignments for reads from each species. The reads were screened in AliView v.1.30 (Larsson 2014) to identify potential chimeras, which were removed from the alignments. Then, all five alignments were split into subalignments by primer and polymerase combination, retaining up to 100 reads per subalignment. In all subalignments, indels and substitutions relative to the consensus sequence were counted. The indel and substitution rates were expressed as per cent per base, accounting for the number of reads in the retained alignment and their modal length. We expected that most of the substitutions reflect polymerase errors, whereas most indels represent sequencing errors (Potapov & Ong 2017).

*De novo* chimera scores were calculated per sample using the UCHIME algorithm (Edgar et al. 2011), implemented in VSEARCH, based on abundance-annotated OTU representatives. Reference-based chimeras were evaluated with VSEARCH against EUKARYOME v2.0. Where relevant, data quality assessments were performed separately for the ultra-long subset and full-ITS data subsets to secure fair comparisons among methods.

In the full-ITS data subset, All OTUs with ITS sequence similarity >99.5% to a reference sequence over >97% of the sequence length were considered of high-quality and non-chimeric. OTUs with total abundance <10 reads were considered of low-quality when their representative sequence had a Phred score <25. OTUs were also assigned as low-quality when they had >2 gap openings compared to substitutions relative to the two best-matching taxa. OTUs with total abundance <3 reads and reference-based or *de novo* chimera score >2 were assigned as chimeric. Singleton and doubleton OTUs with chimera scores >1 were also assigned as chimeric. We further considered OTUs chimeric when they had >99% sequence similarity but 45%-95% coverage relative to the best match, or >98% sequence similarity but 55-90% coverage, or >97% sequence similarity but 55-80% coverage. The rate of index-switching was evaluated using UNCROSS2 (default values; Edgar 2018) and based on the occurrence of mock community members in soil data and vice versa.

In the ultra-long data subset, we first excluded short, incomplete sequences of <2600 bases (Microsporidia lacking the ITS region had a >2608-base marker region in our dataset). Second, we tagged the OTUs that best matched Archaea as non-target organisms. Third, we incorporated the quality assignments from the full-ITS data subset when the OTU representative sequences matched, and we performed the quality evaluation as described above, with additional steps. Additional low-quality reads were assigned when the SSU had >2 gap openings more than substitutions, relative to the two best-matching taxa. Fourth, chimeras were determined for all OTUs with a total abundance of <10 reads and reference-based or de novo chimera score >10.0. The higher threshold was used because the average chimera scores were positively correlated with overall read length. Additional chimeras were manually identified by comparing kingdom-level to order-level taxonomic assignments across the first 500 bases of SSU, the ITS region and the last 500 bases of LSU, with no OTU abundance threshold.

Of the 12,861,972 ultra-long HiFi reads, 564,694 were assigned to soil samples, 81,858 to the 103-species mock community and 135,405 to the 15-species mock community. Across these data, we estimated the overall read proportions for index-switch at 0.008%, chimeras at 4.8%, low-quality at 1.6% and incompleteness at 13.2%.

### 2.5. Statistical analyses

Given the asymmetric design (Figure 1), the data were analysed in five subsets: i) ultra-long-read soil data including five polymerases and two primer pairs (Euk575F-21Rngs and Euk002F-Euk5493R; sample G5917 removed due to poor performance) to assess polymerase effects; ii) full-ITS soil data produced using the Solis polymerase and four primer pairs (including ITS9mun-ITS4ngsUni published previously) to test the primer effects; iii) ultra-long-read 103-species mock community; iv) ultra-long-read 15-species mock community and v) full-ITS 103-species mock community.

In relevant datasets, we evaluated the polymerase and primer effects on the following variables: proportions of index-switched, short, archaeal (i.e., non-target), low-quality and chimeric reads; proportions of substitutions and indels; average OTU read length per sample; proportions of reads belonging to major kingdoms and phyla and Microsporidia (treated as a subphylum within Rozellomycota); and taxonomic richness at the level of OTUs, genera and class-level lineages (including those assigned to kingdom or phylum level only). All richness values were analyzed as residuals from a regression of the logarithm of richness ∼ the logarithm of sequencing depth, to account for differences in sequencing depth and avoid information loss. For illustrative purposes, we also calculated residuals from the regression of richness ∼ logarithm of sequencing depth, with taxon as the unit. Generalized mixed models were used to assess the effects of polymerases, primer pairs and their interaction term, as well as the biological sample (random blocking effect). Spearman correlation coefficients among treatments were calculated from pooled biological samples and logarithm-transformed abundances to reduce the influence of the most common species. Potential read length bias among polymerases and primers was additionally assessed by correlating read abundances of OTUs to the OTU read length in each sample or pooled biological samples, with or without including unobserved OTUs (Tedersoo et al. 2026b).

## 3. Results

### 3.1. Polymerase biases

The selected high-processivity polymerases differed in average sequencing error rates, chimeric read formation and the production of low-quality and partial reads (Figure 3), but not in the average amplicon length of OTUs or index-switching. In the mock community members, substitutions prevailed over indels, suggesting that most errors were introduced during amplification. The substitution rate was greatest in TaKaRa (average, 0.099 errors per base) and Solis (0.068), which were significantly higher than for Fusion (0.014), Platinum (0.008) and Phusion (0.006). The indel rate in TaKaRa (0.050) was higher than that of other polymerases (0.009-0.016), suggesting that most of its indels are PCR-related rather than sequencing artefacts.

**Figure 3.**
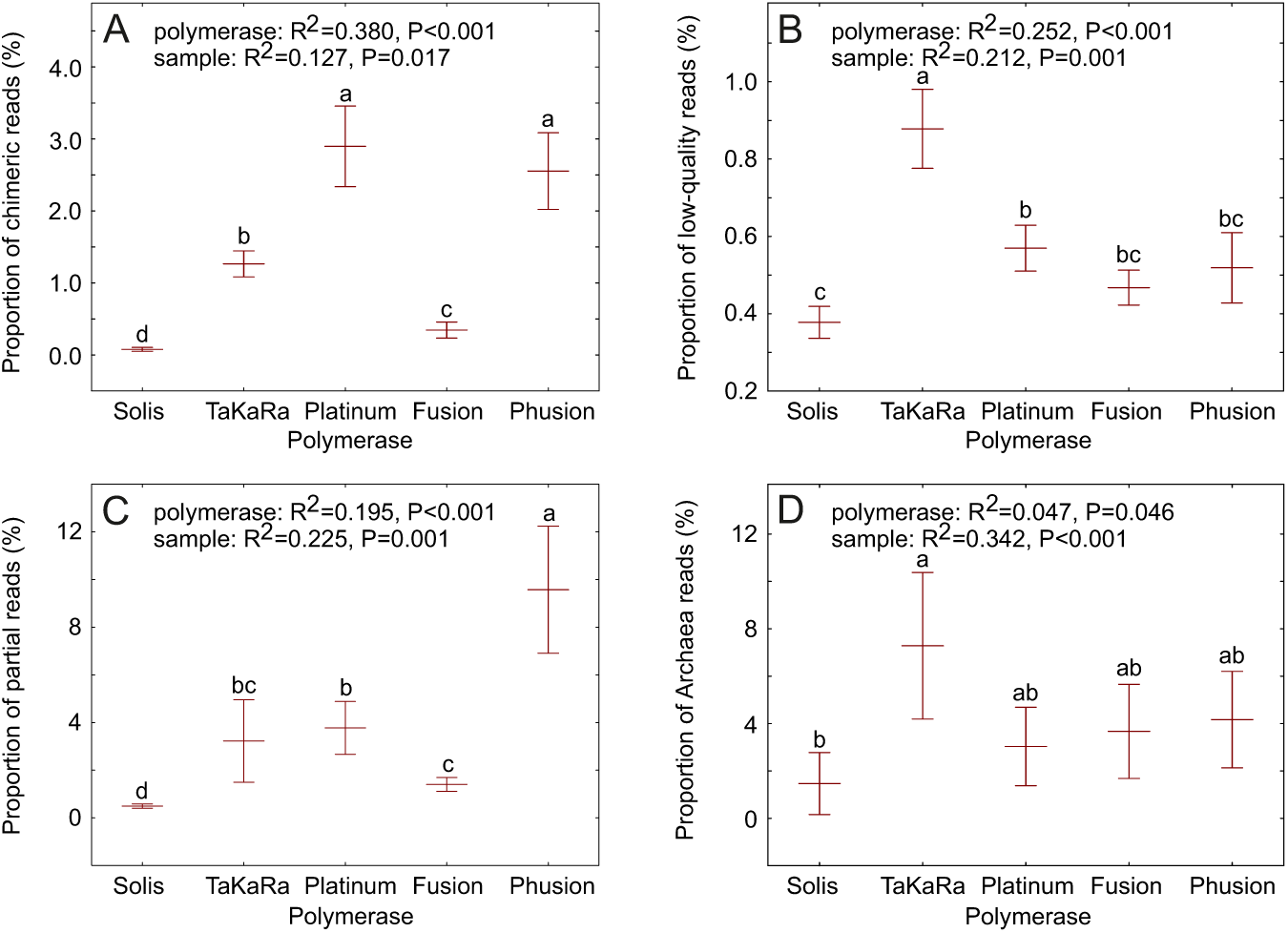
Differences in the performance of five PCR polymerase kits, based on the ultra-long data subset, in A) proportion of chimeric reads, B) proportion of low-quality reads, C) proportion of partial reads and D) proportion of non-target reads assigned to Archaea. Lines and whiskers denote means ± SE; letters above whiskers indicate statistically significantly different groups based on Tukey HSD tests. The initial tests included a biological sample as a random blocking factor.

Based on the soil ultra-long data subset, the Platinum and Phusion polymerases had a higher proportion of chimeric reads (2.9% and 2.5%, respectively) than TaKaRa (1.3%), Fusion (0.3%) and Solis (0.1%). TaKaRa produced the highest proportion of low-quality reads (0.9%), followed by Platinum (0.6%), Phusion (0.5%), Fusion (0.5%) and Solis (0.4%). The proportion of partial reads was highest for Phusion (10.0%), followed by Platinum (3.8%), TaKaRa (3.3%), Fusion (1.4%) and Solis (0.4%). The highest proportion of Archaea reads was produced by TaKaRa (7.3%), followed by Phusion (4.2%), Fusion (3.8%), Platinum (3.0%) and Solis (1.6%).

Platinum and Fusion polymerases recovered significantly more OTUs and classes compared with TaKaRa, Solis and Phusion polymerases in the soil community (Figure 4) but not in the two mock communities of fungi. Phusion, Platinum and Solis recovered more genera compared with TaKaRa and Phusion (Figure 4). Polymerases recovered somewhat different communities at the highest taxonomic levels (Figure 4). Solis and Fusion recovered a higher proportion of Amoebozoa, Fungi, Nematoda, Rhizaria and Stramenopila than TaKaRa, Platinum and particularly Phusion. Platinum recovered the highest proportion of Viridiplantae, followed by Solis, Phusion and Fusion, and TaKaRa distinctly the lowest.

**Figure 4.**
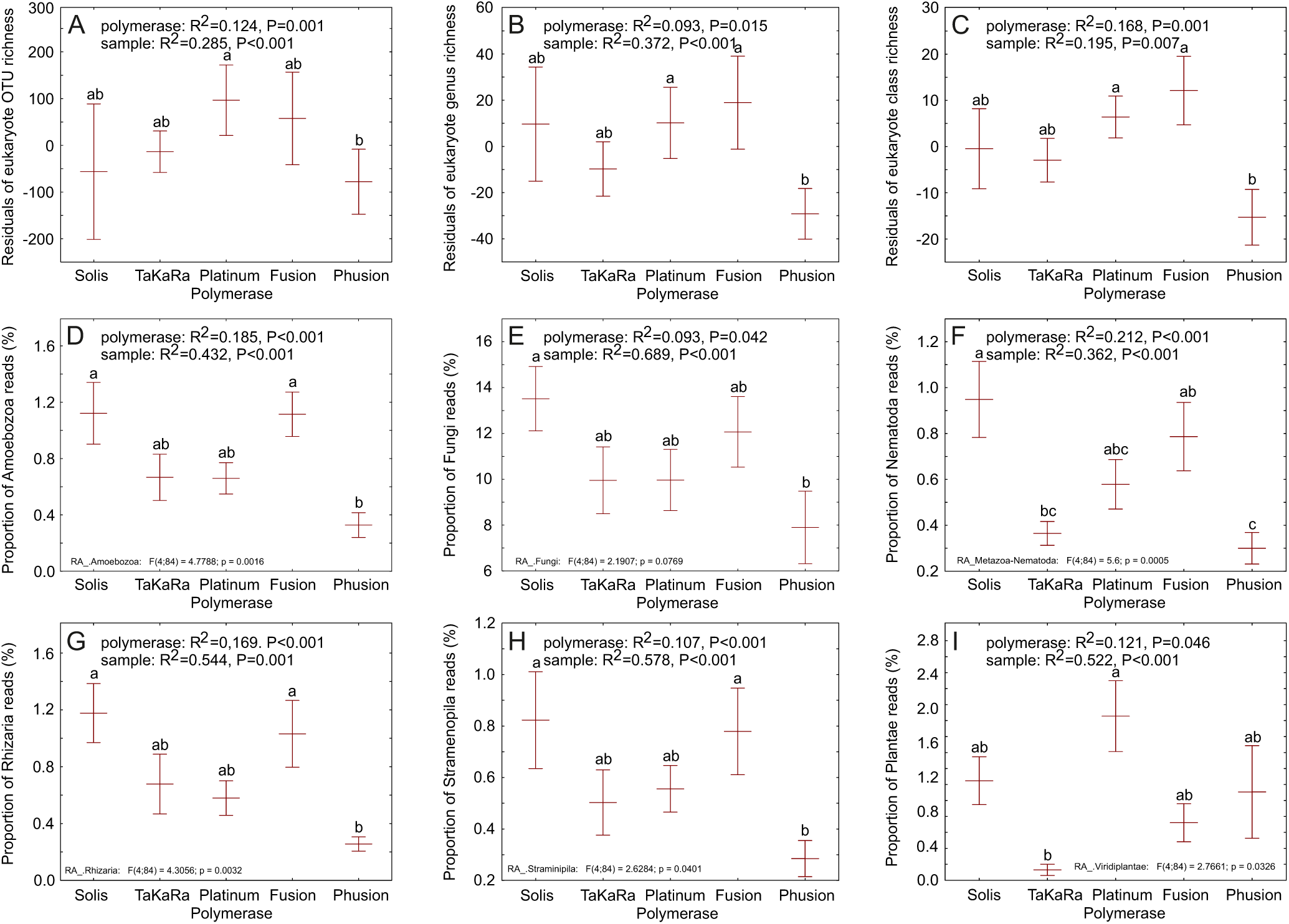
Differences among the performance of five PCR polymerase kits in the recovery of taxonomic richness and major eukaryotic kingdoms and phyla based on the ultra-long data subset: A) OTU richness, B) genus richness, C) class-level lineage richness; D) proportion of Amoebozoa reads; E) proportion of Fungi reads; F) proportion of Nematoda reads; G) proportion of Rhizaria reads; H) proportion of Stramenopila reads; and I) proportion of Plantae reads. Lines and whiskers denote means ± SE; letters above whiskers indicate statistically significantly different groups based on Tukey HSD tests. The initial tests included a biological sample as a random blocking factor.

### 3.2. Primer biases

Based on the Solis full-ITS data subset of soil amplicons, the tested primers differed in the proportions of chimeric reads, low-quality reads, partial reads and Archaeal reads (Figure 5) but not in sequencing error rates, average amplicon length of OTUs or index-switches. Contrary to expectations, the shortest amplicon of ITS9mun-ITS4ngsUni revealed the highest proportion of chimeric reads (mean ± SD, 1.66 ± 2.39%), followed by 3NDF-21R (0.14 ± 0.17%), Euk575F-21Rngs (0.08 ± 0.07%) and the longest amplicon of Euk002F-Euk5493R (0.02 ± 0.03%) within the ITS region. For the ultra-long fragment, the chimeric read proportions increased dramatically for the 3NDF-21R (1.52 ± 0.56%) primer pair but less for the Euk575F-21Rngs (0.16 ± 0.11%) and Euk002F-Euk5493R (0.10 ± 0.05%) primer pairs. Based on the ITS region alone, the ITS9mun-ITS4ngsUni and 3NDF-21R primer pairs produced the highest proportion of low-quality reads, 6.44 ± 3.13% and 2.36 ± 1.45%, respectively. In the ultra-long data subset, the Euk575F-21Rngs primer pair recovered a higher proportion of Archaea reads (2.94 ± 7.79%) than 3NDF-21R (1.06 ± 2.41%) and Euk002F-Euk5493R (0.00 ± 0.00%) primer pairs (F_2,16_=1.25; P=0.030).

**Figure 5.**
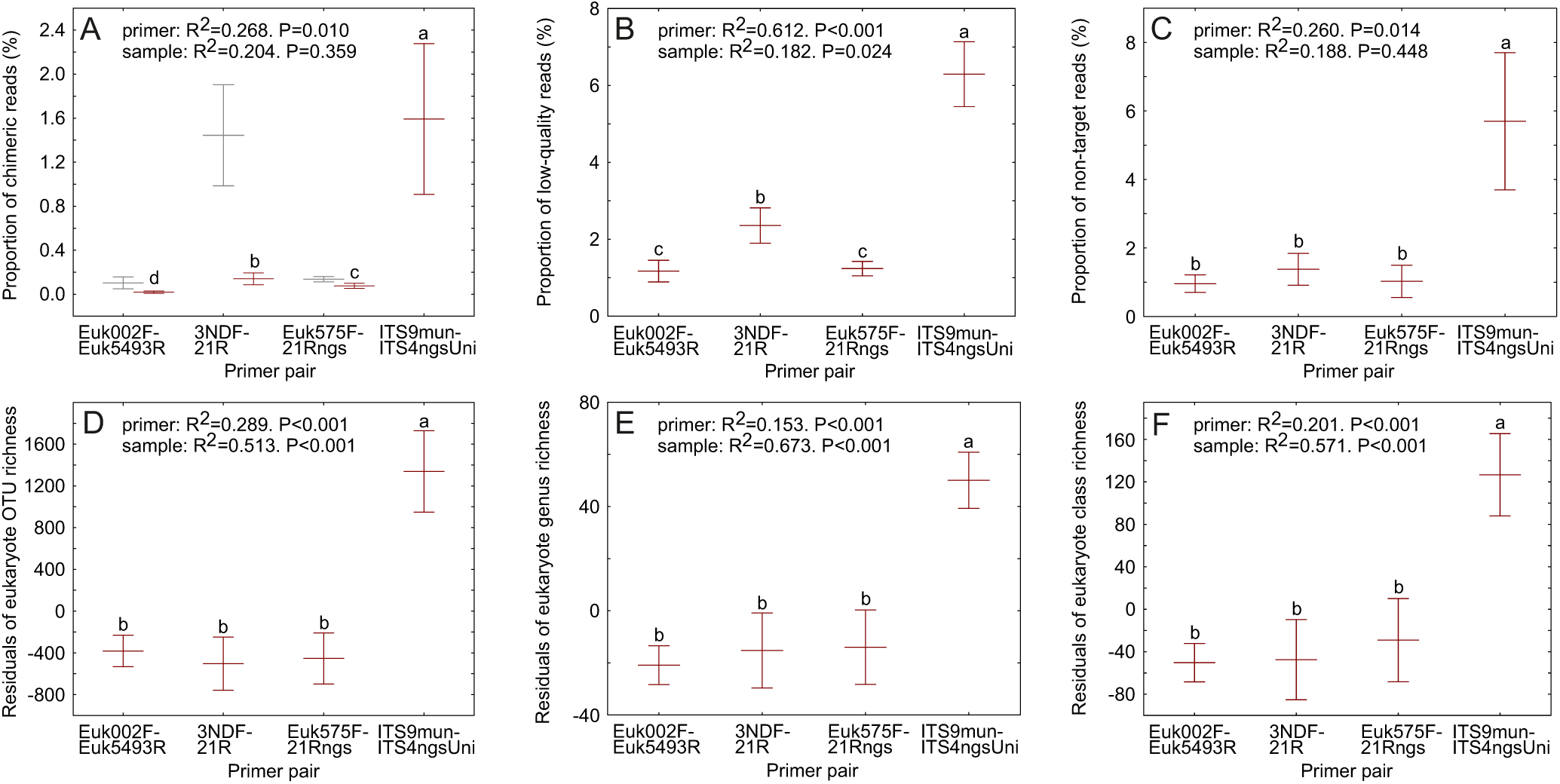
Differences in the performance of four primer pairs in producing technical artefacts and taxonomic richness in the full-ITS data subset: A) proportion of chimeric reads (grey symbols refer to the ultra-long data subset); B) proportion of low-quality reads; C) proportion of non-target reads (including Archaea and metagenomic reads); D) OTU richness; E) genus richness; and F) class-level lineage richness. Lines and whiskers denote means ± SE; letters above whiskers indicate statistically significantly different groups based on Tukey HSD tests. The initial tests included a biological sample as a random blocking factor.

The primers differed greatly in alpha diversity estimates. In the 103-species mock community, the ITS9mun-ITS4ngsUni amplicon recovered 97 species, whereas the ultra-long amplicons recovered 54 to 71 species and collectively 86 species based on the analysis of full-ITS. In the 15-species mock community, ultra-long amplicons revealed 9-12 species (13 species collectively), with no differences among polymerases or primers. In the soil community, the ITS9mun-ITS4ngsUni primer pair recovered on average 1574-1789 more OTUs, 156-177 more genera and 64-71 more class-level taxa per sample compared with the ultra-long amplicons based on the ITS region (Figure 5). There were no differences in taxon recovery among the three primer pairs producing ultra-long reads.

The primer pairs exhibited substantial differences in the higher taxonomic-level composition of soil samples (Figure 6). In general, the full-length ITS primer pair differed the strongest from the ultra-long amplicon primers. The full-length ITS primers overyielded OTUs belonging to Dinoflagellata, Choanoflagellata, Euglenozoa, Fungi (in particular the phyla Glomeromycota, Monoblepharomycota, Rozellomycota and Zoopagomycota) and Tracheophyta (Viridiplantae), but revealed lower proportions of Alveolata, Amoebozoa, Microsporidia, Rotifera and Cercozoa (Figure 6). Of ultra-long amplicon primer pairs, EUK002F-Euk5493F revealed higher proportions of Mucoromycota and Amoebozoa but less Rotifera, Zoopagomycota and Microsporidea. EUK575F-21Rngs yielded higher Choanoflagellata, Rotifera, Zoopagomycota, Monoblepharomycota, Glomeromycota, Dinoflagellata but lower Rozellomycota and Apicomplexa proportions. The 3NDF-21R primer pair revealed lower proportions of Cercozoa, Glomeromycota, Amoebozoa and Dinoflagellata compared with other primer pairs (Figure 6).

**Figure 6.**
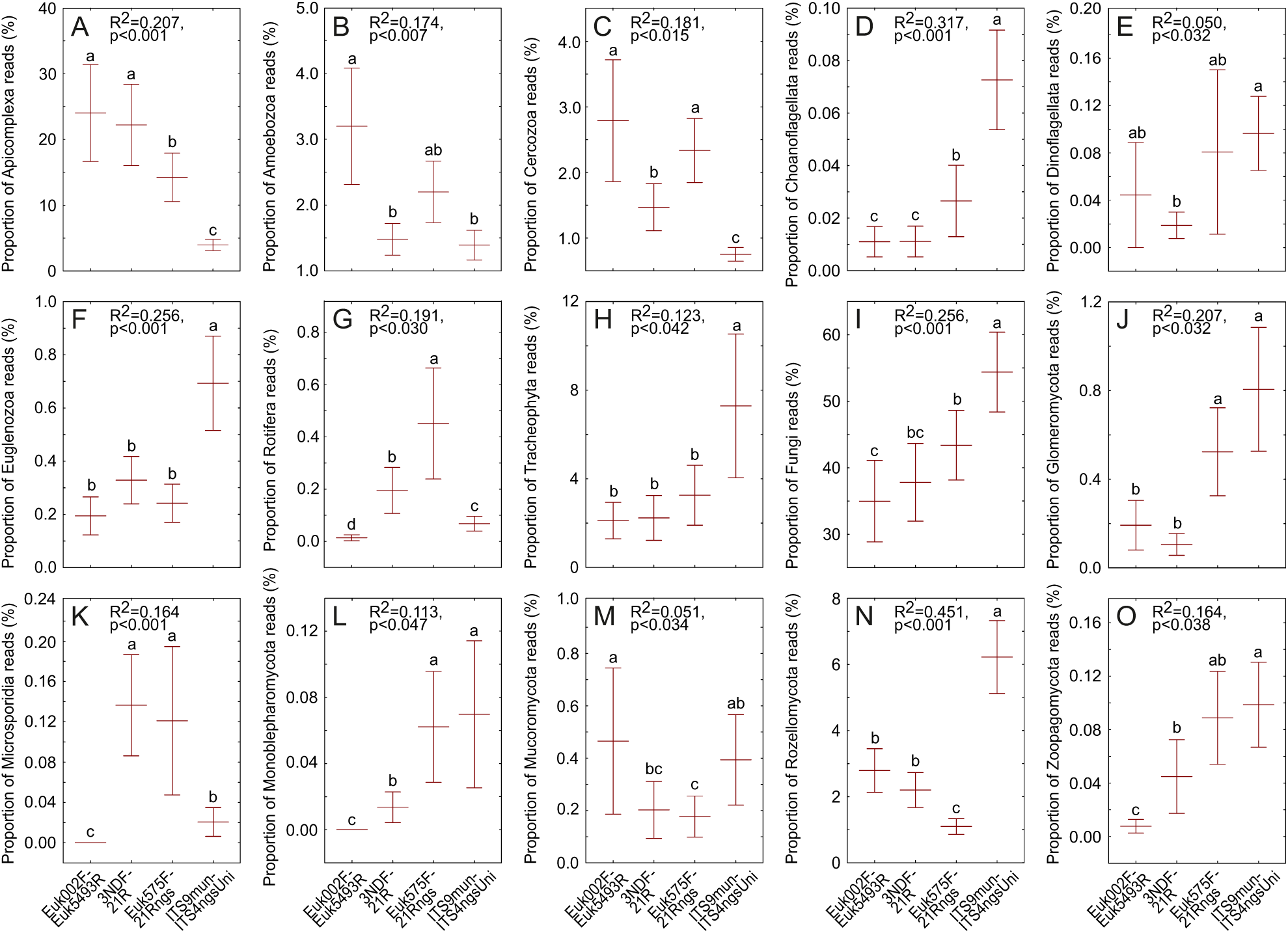
The effect of primer pairs on the relative abundance of major eukaryote kingdoms and phyla based on the Solis full-ITS data subset of soil amplicons: A) proportion of Apicomplexa reads; B) proportion of Amoebozoa reads; C) proportion of Cercozoa reads; D) proportion of Choanoflagellata reads; E) proportion of Dinoflagellata reads; F) proportion of Euglenozoa reads; G) proportion of Rotifera reads; H) proportion of Tracheophyta reads; I) proportion of Fungi reads; J) proportion of Glomeromycota reads; K) proportion of Microsporidia reads; L) proportion of Monoblepharomycota reads; M) proportion of Mucoromycota reads; N) proportion of Rozellomycota reads; and O) proportion of Zoopagomycota reads. Rozellomycota, as displayed here (panel N), does not include Microsporidia (panel K). Lines and whiskers denote means and standard errors. Letters above bars denote statistically significantly different groups when the sample was used as a random blocking factor.

### 3.3. Correlations among primers and polymerases

In the soil full-ITS data subset, sample-based richness estimates were moderately correlated among Fusion, Platinum, Solis and TaKaRa polymerases for the Euk575F-21Rngs primer pair in OTU richness estimates (average R=0.745; range, 0.587 to 0.894), with slightly higher values at the genus- and class-level. The Phusion polymerase showed no significant correlations with other primer pairs (R<0.333; P>0.1) in all richness comparisons. Correlations among polymerases for the Euk002F-Euk5493R primer pair followed similar trends but were somewhat weaker (OTU richness: average R=0.636; range, 0.386 to 0.744) in all three taxonomic levels. The correlations of richness estimates of the Euk575F-21Rngs and Euk002F-Euk5493R were relatively lower for Phusion (average R=0.595; range, 0.446 to 0.756) and Fusion (average R=0.666; range, 0.543 to 0.783) polymerases than others (R>0.883). A comparison of four primer pairs in the Solis polymerase data subset revealed moderately strong relationships among the three ultra-long amplicon primer pairs for OTU richness (average R=0.749; range, 0.621 to 0.843), genus richness (average R=0.862; range, 0.829 to 0.902) and class-level richness (average R=0.723; R range, 0.581 to 0.796). The ITS9mun-ITS4ngsUni primer pair had weak, non-significant positive correlations with other primer pairs at all taxonomic levels.

Within the ultra-long amplicons, richness comparisons between full-ITS and ultra-long data subsets revealed strong correlations among polymerases at the OTU, genus and class levels. These correlations were strongest for Solis (average R=0.996) and Fusion (average R=0.986), somewhat lower for Platinum (average R=0.956) and Phusion (average R=0.938), and lowest for TaKaRa (average R=0.691). The primer pairs showed no differences in richness correlations within the Solis polymerase data subset (R>0.960).

## 4. Discussion

### 4.1. PCR polymerase kits

Our study indicates that high-processivity polymerases exhibit great differences in generating PCR errors, chimeras and low-quality reads. The widely used TaKaRa polymerase performs the worst, generating high levels of both substitutions and indels during PCR. The Solis polymerase also has relatively high substitution rates compared with Fusion, Phusion and Platinum polymerases. However, Solis produced the lowest proportions of chimeric, low-quality and non-target reads assigned to Archaea. In terms of chimera formation, Platinum, Phusion and TaKaRa performed relatively poorly, corroborating previous studies for the two former polymerases in short-read applications (Bakker 2018; Sze & Schloss 2019). TaKaRa and Phusion also revealed relatively high proportions of partial reads and the non-targeted Archaea with ∼3000-base reads. Their suboptimal performance may be related to amplicon size limits, insufficient DNA quality, or loss of primer specificity under the used thermal cycling conditions or during polymerase proofreading (Gohl et al. 2021).

The selection of polymerases may affect biodiversity recovery (Bakker 2018, Nichols et al. 2018). Here, Platinum and Fusion kits revealed the highest number of eukaryote OTUs. For Platinum, it may be at least partly related to the undiscovered chimeras given its high overall chimera rate. Solis and Fusion polymerases recovered the highest proportions of the best-known kingdoms and phyla, whereas other polymerases yielded more groups with limited taxonomic information and potentially artefactual origin. Polymerases displayed the greatest differences in sequencing plant DNA, which we cannot explain. Short-read studies show that Phusion and Platinum exhibit greater variability than other polymerases in recovering mock community members (Nichols et al. 2018).

The main advantages of the Fusion polymerase are its relatively low chimera-formation and PCR-error rates, but its main disadvantage is its relatively low community correlation with other high-processivity polymerases, suggestive of taxonomic biases. Despite its relatively high substitution rate, the Solis polymerase has the advantage of producing the lowest proportions of partial, chimeric and low-quality reads, the best recovery of sequences from the most common eukaryote lineages and being used in EUKARYOME database-related studies, which enable cross-comparison of thousands of biological samples. Given the high PCR error and chimerism rates and relatively poor biodiversity recovery, we recommend avoiding the use of TaKaRa, Phusion and Platinum polymerases for >4-kb amplicon sequencing.

### 4.2. Primer pairs

Our analyses showed major differences in higher-taxon recovery among the ultra-long amplicon primer pairs, especially for Apicomplexa, Microsporidia and Rotifera, which is only partly explained by primer-template mismatches. Based on in vivo evidence, the Euk575F-21Rngs primer pair discriminated against the fewest taxonomic groups – Rozellomycota – consistent with in silico analyses. This primer pair had the strongest average correlations to other primer pairs, including ITS9mun-ITS4ngsUni. The relatively low proportions of low-quality and partial reads, high taxonomic coverage and standard use in the EUKARYOME database project render the Euk575F-21Rngs primer pair a good choice for ultra-long-read analyses of soil eukaryote communities. However, given the relatively high recovery of Archaea, these primers may be suboptimal for anoxic sediments or when combined with TaKaRa or Phusion polymerases. The Euk002F-Euk5493R primer pair may be more suitable in such Archaea-rich environments and in situations where nearly the entire SSU and LSU genes are required.

In previous studies, primer selection and rRNA marker regions revealed major differences in the recovery of various eukaryotic groups (Tedersoo et al. 2015; de Filippis et al. 2017; Stuart et al. 2024). Tedersoo & Anslan (2019) reported that the ITS9mun and ITS4ngsUni primers are the least biased among eukaryote full-ITS primers. Here, the ITS9mun-ITS4ngsUni amplicon generally exhibited greater differences to ultra-long reads compared to the among-primer-pair differences in ultra-long amplicons. Of taxonomic groups, the ITS9mun-ITS4ngsUni primer pair recovered relatively more Dinoflagellata, Choanoflagellata, Euglenozoa, Tracheophyta and various fungal phyla but relatively less Apicomplexa, Amoebozoa, Rotifera, Cercozoa and Microsporidia than the ultra-long-read primers. Both the over- and underrepresented groups included taxa with relatively short (e.g., Apicomplexa, Microsporidia and Euglenozoa) and long (e.g., Amoebozoa, Choanoflagellata and Cercozoa) marker regions, which is consistent with other analyses showing that there is negligible read-length bias for the PacBio sequencing platform in the 700-6000 bp amplicon size range (Tedersoo et al. 2026b). Studies using the Illumina sequencing platform have found that strong marker and primer effects are associated with GC content and marker length (Shaffer et al. 2025).

The ultra-long primer pairs differed in the read quality. The 3NDF-21R primer pair yielded more chimeric and low-quality reads than other primer pairs tested. The higher proportion of chimeric reads of the 3NDF-21R primer pair increased specifically when considering chimeras in the ultra-long fragment compared with the full-ITS fragment. Unexpectedly, the shortest ITS9mun-ITS4ngsUni amplicon yielded a greater proportion of low-quality and chimeric reads compared with ultra-long amplicons despite greater circular consensus coverage and less potential chimera breakpoints. However, this may be related to differences in the type of Solis polymerase used for producing ultra-long and full-ITS amplicons.

### 4.3. Ultra-long and full-ITS amplicon comparisons

The among-study comparison revealed that using the same DNA extract, full-ITS primers recover 40%-110% higher richness than ultra-long-read primers in the mock community, documenting 94% of the input species. We speculate that the relatively poor performance of ultra-long amplicons in the mock community may be related to low DNA quality of some of the pooled fruit-bodies rather than primer or polymerase biases, because the collective species recovery of multiple ultra-long amplicons was still 89% of the full-ITS amplicon.

Furthermore, the ITS9mun-ITS4ngsUni primer pair recovered 3-fold to 9-fold more OTUs than the ultra-long primer pairs in the soil community obtained from the same DNA extracts based on the full-ITS data subset. A part of this may be explained by higher proportions of various artefacts in the full-ITS amplicon, many of which may have been overlooked. Besides OTU counts, a better performance of the ITS9mun-ITS4ngsUni primer pair is reflected in 106%-352% more genera and 58%-175% more class-level lineages recorded. In a preprint, Dai et al. (2026) suggested that 0.4-kb SSU markers are similarly more efficient than 1.8-kb markers in biodiversity monitoring of aquatic protists. Collectively, these observations point to potential issues with DNA quality, as long markers are less likely to be amplified from fragmented DNA. Furthermore, weak correlations in richness among primer pairs suggest that ultra-long amplicons may be unsuitable for comparisons of richness and biodiversity in soil eukaryote communities.

Although ultra-long reads reach 3-6 kb, their sequence-similarity-based taxonomic identification typically focuses on a single subregion, such as SSU V4-V5 or ITS. This is mainly attributable to the paucity of full-length reference sequences and to sporadic coverage of the marker subregions, which differ across taxonomic groups. A vast majority of reference sequences of cultures or herbarium specimens have been produced using Sanger sequencing of amplicons <1000 bp. It would require bidirectional Sanger sequencing of 5-8 amplicons to generate nearly full-length SSU-ITS-LSU sequences without gaps - i.e., a substantial effort considering the resources. Apart from phylogenetic placement algorithms that may fail to handle large, non-overlapping regions, no analytical pipelines are available to simultaneously account for different subregions or to automatically select marker information most relevant to a particular taxon. Similarly, our analysis required manual accounting for SSU and LSU beyond the main ITS region, which took extra time and reduced reproducibility. These analyses implied that the LSU gene provided little additional taxonomic information due to high redundancy with ITS and typically confirmed taxonomic identity at the genus level or higher. Conversely, the SSU V4-V5 subregion constituted an important additional source for genus- to class-level taxonomic assignments in many animal phyla but much less in protists. However, the V1-V3 subregion of SSU is broadly used in certain protists, such as amoebozoans and excavates, but two of the three ultra-long marker primer pairs did not cover this part of SSU.

For the ultra-long amplicon analysis of the 103-species mock community, the trimmed full-ITS OTUs were more informative than the ultra-long-read OTUs. This is related to the fact that ultra-long reads included tens to hundreds of rare variants in common species, representing undetected chimeras and reads with single, long indels (5-20 bases) in SSU or LSU. Because of these multiple artefacts and high sequence similarity among congeneric *Cortinarius* species, the 103-species mock community was impossible to evaluate accurately for the ultra-long barcode. Conversely, the full-ITS subset was straightforward to assess due to fewer long-indel artefacts and higher average taxonomic resolution resulting from trimming off the SSU and LSU genes uninformative at the species level. Nonetheless, the taxonomic richness estimates based on full-ITS and ultra-long-read OTUs were strongly correlated in soil samples (R > 0.95).

## 5. Conclusions and perspectives

Our study indicates that the selection of polymerases and primer pairs and marker size all affect the recovery of soil eukaryote biodiversity. Compared with other tested polymerases, the Fusion and Solis polymerases yield relatively robust results despite trade-offs in error rates. Alarmingly, the ultra-long amplicon primer pairs show marked differences in the recovery of major eukaryote groups, underscoring the importance of this information for meta-studies. The ultra-long amplicons, preferably generated with the Euk575F-21Rngs primer pair, are essential for testing community-level phylogenetic hypotheses and for manually or semi-automatically identifying taxa as accurately as possible. Although ultra-long reads offer great potential for evolutionary and ecological analyses (Jamy et al. 2022) and the description of previously unrecognized lineages (Tedersoo et al. 2026a), their proper utilization lags far behind in bioinformatics pipelines. Such analytical methods require urgent development to leverage the broad spectrum of taxonomic and phylogenetic information that ultra-long reads provide.

Shorter amplicons recover many more OTUs and higher taxa than ultra-long amplicons, suggesting the preferential use of the ITS9mun-ITS4ngsUni primer pair or other universal primer pairs targeting medium-size DNA fragments for surveys documenting and comparing soil eukaryote biodiversity. It remains to be tested to what extent the marker length is related to co-amplification and sequencing of extracellular DNA.

## Author Contributions

L.T. designed research, analysed data and wrote the paper, with contributions from all co-authors; V.F.A. calculated error rates based on mock communities; R.P. performed laboratory analyses; V.M. performed bioinformatics analyses.

## Acknowledgements

We thank Irma Zettur and Irja Saar for compiling the mock communities. We acknowledge the Norwegian Sequencing Centre (Oslo) for the PacBio sequencing services.

## Data Availability Statement

The raw sequencing reads generated in this study have been deposited in the European Nucleotide Archive (ENA) under project accession PRJEB123023 (sample accession numbers SAMEA123257319 - SAMEA123257330; sequence accession numbers ERR17666980 - ERR17667127. The Bash and R workflows, together with documentation of the required inputs and processing steps are available in GitHub (https://github.com/Mycology-Microbiology-Center/euk-ultralong-biases) and Zenodo (DOI:10.5281/zenodo.23687374).

## Benefits Sharing Statement

Benefits from this research accrue from the sharing of our data and results on public databases as described above.

## Ethics statement

Our work follows all ethical and research integrity standards.

